# Brain tissue segmentation based on MP2RAGE multi-contrast images in 7 T MRI

**DOI:** 10.1101/455576

**Authors:** Uk-Su Choi, Hirokazu Kawaguchi, Yuichiro Matsuoka, Tobias Kober, Ikuhiro Kida

## Abstract

We proposed a method for segmentation of brain tissues––gray matter (GM), white matter (WM), and cerebrospinal fluid (CSF)—using multi-contrast images, including a T1 map and a uniform T1-weighted image, from a magnetization-prepared 2 rapid acquisition gradient echoes (MP2RAGE) sequence at 7 Tesla. The proposed method was evaluated with respect to the processing time and the similarity of the segmented masks of brain tissues with those obtained using FreeSurfer, FSL, and SPM12. The processing time of the proposed method (28 ± 0 s) was significantly shorter than those of FSL and SPM12 (444 ± 4 s and 159 ± 2 s for FSL and SPM12, respectively). In the similarity assessment, the tissue mask of the brain obtained by the proposed method showed higher consistency with those obtained by FSL than with those obtained by SPM12. The proposed method misclassified the subcortical structures and large vessels since it is based on the intensities of multi-contrast images obtained using MP2RAGE, which uses a similar segmentation approach as FSL but is not based on a template image or a parcellated brain atlas, which are used for FreeSurfer and SPM12, respectively. However, the proposed method showed good segmentation in the cerebellum and WM in the medial part of the brain in comparison with the other methods. Thus, because the proposed method using different contrast images of MP2RAGE sequence showed the shortest processing time and similar segmentation ability as the other methods, it may be useful for both neuroimaging research and clinical diagnosis.

## Introduction

Structural information regarding brain tissue is important for both neuroimaging research and clinical diagnosis. Magnetic resonance imaging (MRI) has been widely used to obtain structural information from various types of contrast images. Different MR contrast images can show brain abnormalities via segmentation of subcortical structures in neuronal disorders, such as Parkinson’s or Alzheimer’s diseases [1]. Furthermore, gray matter (GM) segmentation can be used to estimate cortical thickness or volume to evaluate developmental stages or the effects of aging [2]. In functional MRI, white matter (WM) segmentation can provide an inflated brain mesh [3] to project brain activation maps.

Most previous MRI segmentation methods for brain tissues, including GM, WM, and cerebrospinal fluid (CSF), were based on the signal intensities in T1-weighted (T1w), T2-weighted (T2w), and proton-density (PD) images. However, these images show an intrinsic overlapping intensity distribution among brain tissues, which renders their segmentation into GM, WM, and CSF challenging. In addition, the overlap can often be substantial because of factors such as image inhomogeneity, noise, and the partial volume effect (PVE), which can reduce the precision of the segmentation. Among these factors, PVE is a critical obstacle in brain tissue segmentation because other artifacts can be improved or corrected with sophisticated pre-processing methods, such as bias field correction [4], but PVE cannot be easily corrected because it occurs as a result of the limited spatial resolution of MRI systems.

To overcome the effects of PVE in brain tissue segmentation, numerous groups have employed several approaches using (1) thresholding methods, (2) region-growing methods, (3) clustering, and (4) Bayesian classification [5–7]. In the thresholding method, brain tissues are defined by thresholds based on their intensity histograms. This approach is fast and efficient but may fail to accurately define brain tissues because it does not consider neighborhood information [8]. The region-growing method considers the information of neighborhood voxels by estimating their similarity to a seed voxel, which can lead to biased results because of incorrect seed point selection. The clustering methods use intensities and spatial information (i.e., neighborhood information) as features and classify brain voxels of different types of brain tissue without requiring training. A previous study increased the classification accuracy using various clustering algorithms, such as expectation–maximization or fussy C-means [9]. Finally, Bayesian classification has been adopted by several popular segmentation software packages, such as SPM12, FSL, and FreeSurfer [10]. Zhang et al. (2001) calculated the maximum probabilities of each brain tissue and used the neighborhood voxel information by employing the Markov random field statistical model to improve brain tissue classification [11]. However, these intensity-based approaches still involve PVE MRI artifacts, particularly in the segmentation of infant brains [12] or abnormal brain structures, such as a tumor [13]. Machine-learning algorithms have also been applied to brain segmentation methods [14].

Recently, MRI systems have shifted to using ultra-high fields [UHF ≥ 7 Tesla (T)] to obtain fine-structure and functional images with sub-millimeter resolution. However, acquisition of total brain volume images at sub-millimeter resolutions increases the dataset size substantially, and the segmentation methods described above require longer computation times for such high spatial-resolution images. In addition, T1w images for brain tissue segmentations are usually obtained with a magnetization-prepared rapid gradient echo (MPRAGE) sequence, but the image may contain severe inhomogeneities at UHF. In contrast to the MPRAGE sequence, the magnetization-prepared 2 rapid acquisition gradient echoes (MP2RAGE) sequence can produce a less inhomogeneous T1w image, termed a uniform (UNI) image, by combining two different gradient echo images with two different inversion times (TIs) [15–17]. The MP2RAGE sequence also produces images with different types of contrast, such as two gradient echo images (INV1 and INV2), with different TIs and flip angles (FAs), a T1 map, and a T1w image without a noisy background (UNIDEN).

In the present study, we proposed a segmentation method for brain tissue (GM, WM, and CSF) that used the different contrast images (INV1, INV2, UNI, and a T1 map) acquired by MP2RAGE sequences at 7 T. We evaluated the proposed method with respect to processing time and the similarity of the segmented masks of brain tissues with those from FreeSurfer, FSL, and SPM12, which are commonly used in neuroimaging protocols.

## Methods

### Subjects

Seven volunteers (four males and three females, aged 23–63 years) without a history of neurological disease or any other medical condition participated in this study after providing written informed consent. All experiments were approved by the Ethics and Safety Committees.

### MRI acquisition

The experiments were performed on a 7-T investigational MRI scanner (MAGNETOM 7T; Siemens Healthineers, Germany) with a 32-channel head coil (Nova Medical, Wilmington, MA). The MP2RAGE sequence [15] was acquired using a work-in-progress software package from Siemens Healthineers and using the following parameters: repetition time = 5000 ms, echo time = 3.36 ms, TI1/TI2 = 800 ms/2600 ms, FA1/FA2 = 4°/5°, matrix = 320 x 320 × 256, voxel size = 0.8 × 0.8 × 0.8 mm^3^, integrated parallel acquisition techniques with parallel imaging acceleration = 3, and scan time = 8 min 23 s.

### Prerequisites: brain extraction

The skin and skull from all MP2RAGE images were stripped to extract the brain using the BET toolbox in FSL 5.0 (FMRIB, Oxford, UK). Most images except for UNIDEN could not be automatically stripped because of their different contrasts in comparison with the conventional T1w image and the background noise in these images, such as the salt-and-pepper noise found in UNI. Therefore, the brain-extracted INV2 was used as a mask for the brain extraction process in the other images (UNI, T1, and INV1). After the INV2 mask-based brain extraction, an erosion process was used to remove residual non-brain tissues.

### FreeSurfer segmentation

Brain-extracted UNI images from all subjects were segmented using the auto-reconstruction processes in FreeSurfer 6.0 (https://surfer.nmr.mgh.harvard.edu/fswiki). Since skull stripping had already been performed with the BET toolbox in FSL, this step was excluded in the auto-reconstruction processes using FreeSurfer. After auto-reconstruction, we manually created masks of three brain tissues (GM, WM, and CSF) using the entire parcellated image. In addition, the brainstem, pallidum, and ventral diencephalon [18] were defined as WM for comparison with FSL and SPM12 because these areas are considered to be WM in FSL and SPM12.

### FSL segmentation

Brain-extracted UNI images from all subjects were segmented using the FAST toolbox in FSL 5.0 (http://fsl.fmrib.ox.ac.uk/fsl/fslwiki/). Masks of brain tissues (GM, WM, and CSF) were constructed by binarization of probability maps, which were created in the segmentation process, with a threshold value of 0.5.

### SPM12 segmentation

Brain-extracted UNI images from all subjects were segmented by a unified algorithm that included bias correction, tissue classification, and registration using SPM12 (http://www.fil.ion.ucl.ac.uk/spm/). Probability maps of brain tissues (GM, WM, and CSF) were created using medium bias regularization and a reference brain probability map; then, masks of brain tissues were constructed by binarization of the probability maps with a threshold value of 0.5.

### Image calculation for brain tissue segmentation

A scheme of the proposed method is shown in Fig 1. MP2RAGE images exhibit different contrasts for brain tissues (GM versus WM and CSF versus GM) and different scales of intensity. Therefore, before brain tissue segmentation, we normalized the intensities of the whole brain after stripping the skin and skull for using a feature-scaling method described by Eq. 1. *S_raw_*, min (*S_raw_*), and max (*S_raw_*) represent the raw intensities, the minimum intensity, and the maximum intensity in each MP2RAGE image, respectively. Normalized T1 (nT1), normalized INV1 (nINV1), and normalized UNI (nUNI) contained the normalized intensities.
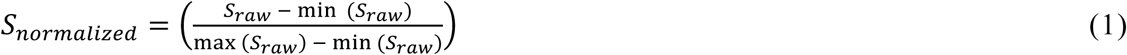

**Fig 1.**
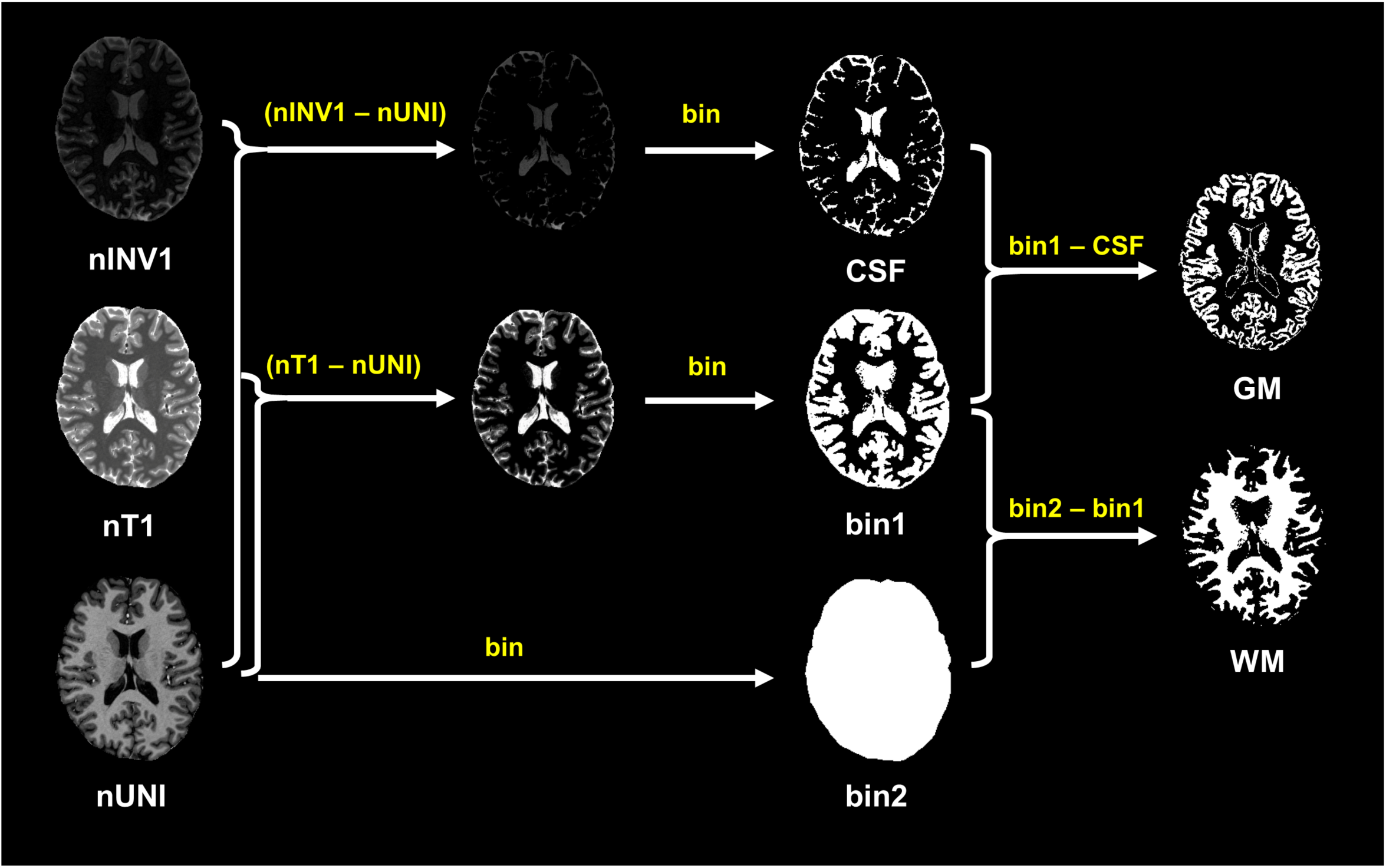
Scheme of the proposed method. nT1: normalized T1, nINV1: normalized INV1, nUNI: normalized UNI, bin: binarization.

We found that each brain tissue produced different normalized intensities based on the different MP2RAGE images. To evaluate the relationship of the intensities between different MP2RAGE images, the intensities of nUNI, nT1, and nINV1 were extracted from all masks of brain tissue segmented by FreeSurfer, FSL, and SPM12 (Fig 2). CSF exhibited larger intensity distributions in the order nT1 > nINV1 > nUNI. GM exhibited larger intensity distributions in the order nT1 > nUNI > nINV1. WM showed larger intensity distributions in the order nUNI > nT1 > nINV1. The calculated values of (nINV1 − nUNI) and (nT1 − nUNI) can clarify the relationships of the normalized intensities between brain tissues (Fig 3). The proposed method is based on the different contrast relationships between simultaneously acquired images. Each mask of brain tissue was calculated using Eqs. 2–4. Binarization with a threshold of 0 is represented by “bin.”
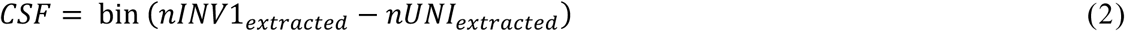

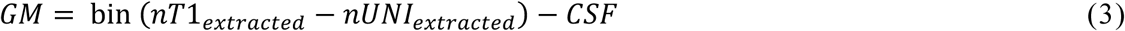

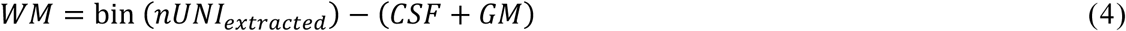

**Fig 2.**
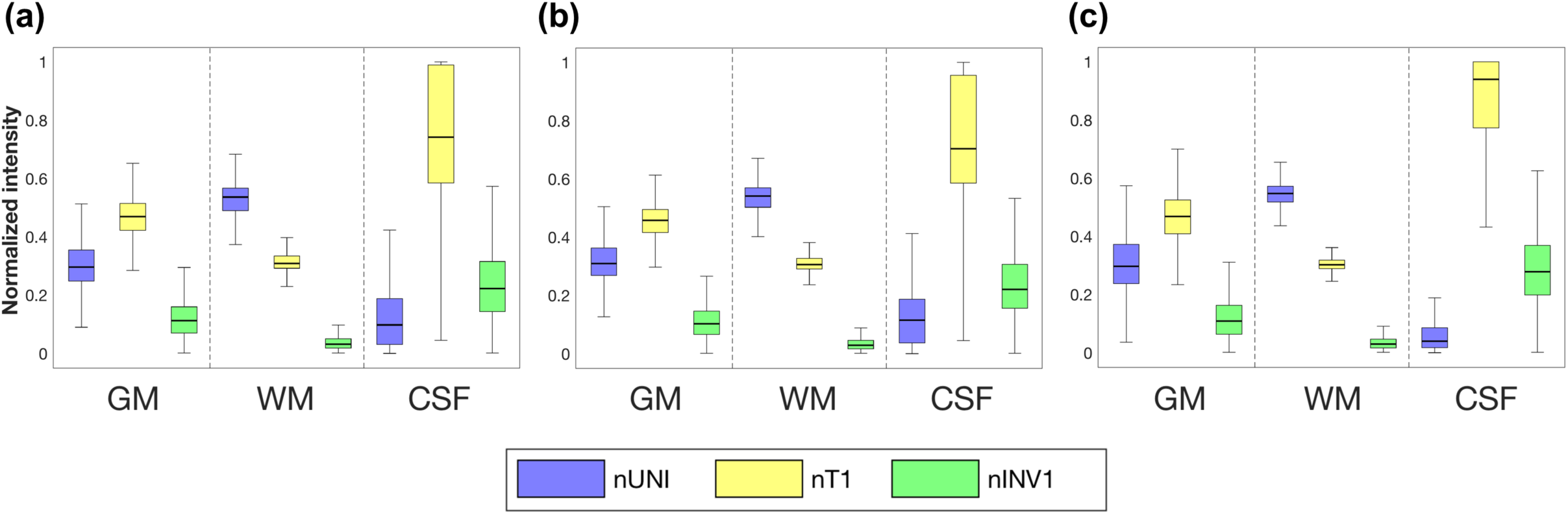
Intensities in brain tissues. Box plots show the intensities of nUNI, nT1, and nINV1 extracted from the tissue masks by FreeSurfer (a), FSL (b), and SPM12 (c).

**Fig 3.**
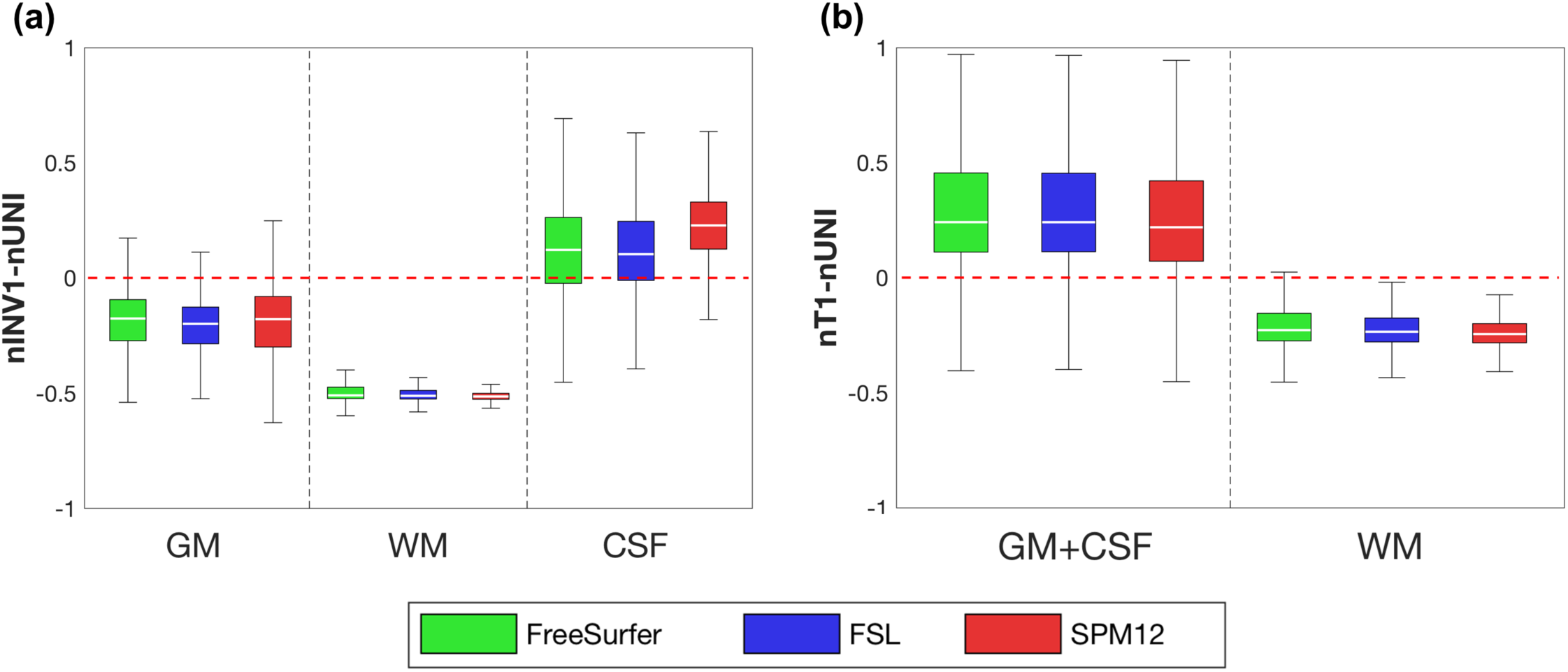
Calculated intensities in brain tissues. Box plots show (nINV1 − nUNI) values (a) and (nT1 − nUNI) values (b) from the FreeSurfer, FSL, and SPM12 masks.

### Evaluation of segmentation

To evaluate the performance of the segmentation, we compared the processing time and the similarity of the masks of brain tissues segmented by the proposed method with those produced with FreeSurfer, FSL, and SPM12. All computations were processed by a computer with the following specifications, and the processing times were recorded: macOS (Sierra 10.12.5), 3.5 GHz 6-Core Intel Xeon E5 CPU, and 16 GB RAM. A paired t-test was performed for comparison of processing times between the proposed method and FSL and SPM12, respectively. For the similarity evaluation, the absolute volume difference (AVD), which measures differences in voxel ratios, and dice coefficient (DICE), which measures the spatial overlap ratio of each mask of brain tissue, were calculated with Eqs. 5 and 6, respectively; the values for these parameters generated by the proposed method and by the other software packages were compared.
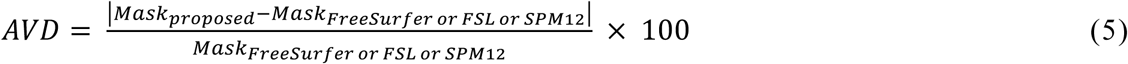

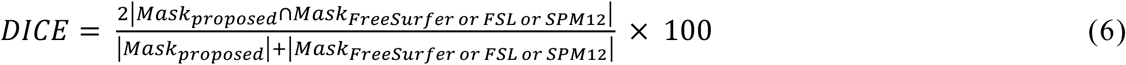

In addition, the modified Hausdorff distance (MHD), which measures boundary distances [19], was calculated using the edges of all tissue masks and compared between the proposed method and the other software packages. The MHD values for each brain tissue were calculated in the same axial slice.

### Estimation of influence of noise on the proposed method

To examine the influence of noise on the proposed method, we added Gaussian noise levels of 3%, 5%, 7%, and 9% to all images and performed the proposed method. The similarity of the masks of brain tissues with noise was evaluated with the masks obtained by FSL without noise as mentioned above.

## Results

### Processing time

The average processing times for the segmentation were 28 ± 0.2 s with the proposed method, 444 ± 4 s with FSL, and 159 ± 2 s with SPM12 (Fig 4). The processing time of the proposed method was significantly shorter than that of the other packages (p < 0.001 vs. FSL and p < 0.001 vs. SPM12, paired *t*-test). The processing time with FreeSurfer was excluded because it included several processing steps beyond brain tissue segmentation, such as the reconstruction of the inflated brain, segmentation of the subcortical structure, and calculation of cortical thickness.

**Fig 4.**
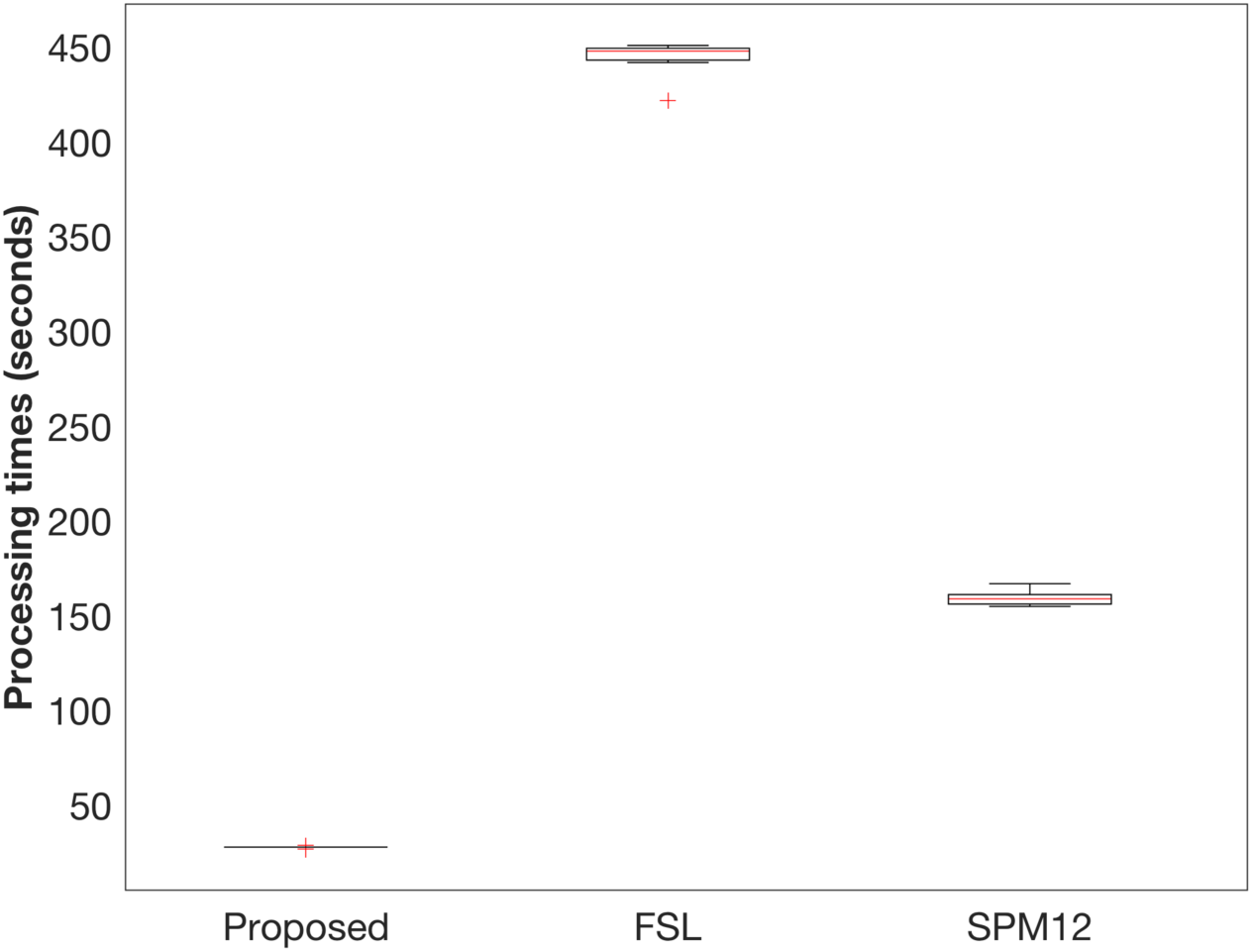
Processing times for the proposed method, FSL, and SPM12. Box plots show the processing times for all subjects with the proposed method (a), FSL (b), and SPM12 (c). The red cross indicates outliers.

### Similarity evaluation

Fig 5 shows the evaluation of the proposed method in comparison with FreeSurfer, FSL, and SPM12 with respect to the AVD, DICE, and MHD of the segmented masks of brain tissues. The spatial differences, i.e., AVD, between the proposed method and FreeSurfer were <20% in all three brain tissues, but the differences were larger (30%–50%) in comparison with SPM12 (Fig 5a). The GM masks segmented by the proposed method almost completely corresponded to those produced by FSL, indicating that AVD was 3.5% ± 1.0%. The average DICEs for GM and WM segmentation between the proposed method and all other methods were >80%, whereas those for CSF segmentation were approximately 75%–82% (Fig 5b). The average DICEs for all brain tissues between the proposed method and FSL were slightly higher than those with FreeSurfer and SPM12. The average MHDs for GM, i.e., 2.3–2.5 mm, between the proposed method and all other methods were larger than those for WM and CSF, i.e., 1.5–1.8 mm (Fig 5c). As for the influence of noise levels, the similarities in AVD, DICE, and MHD in comparison with FSL linearly deteriorated with increasing noise levels in all contrast images of MP2RAGE. However, at a noise level of 9%, the similarities in AVD and DICE were still better than those with SPM12 (Fig 5). Larger differences in segmentation in comparison with SPM12 were found in the GM in the inferior frontal gyrus, cerebellum (Fig 6a), subcortical structures (Fig 6b), large vessels (Fig 6b and 6c), and WM in the medial part of the brain areas (Fig 6c). GM in the inferior frontal gyrus and cerebellum (Fig 6a), and WM in the medial part of the brain areas were well segmented by the proposed method, whereas subcortical structures and large vessels were misclassified.

**Fig 5.**
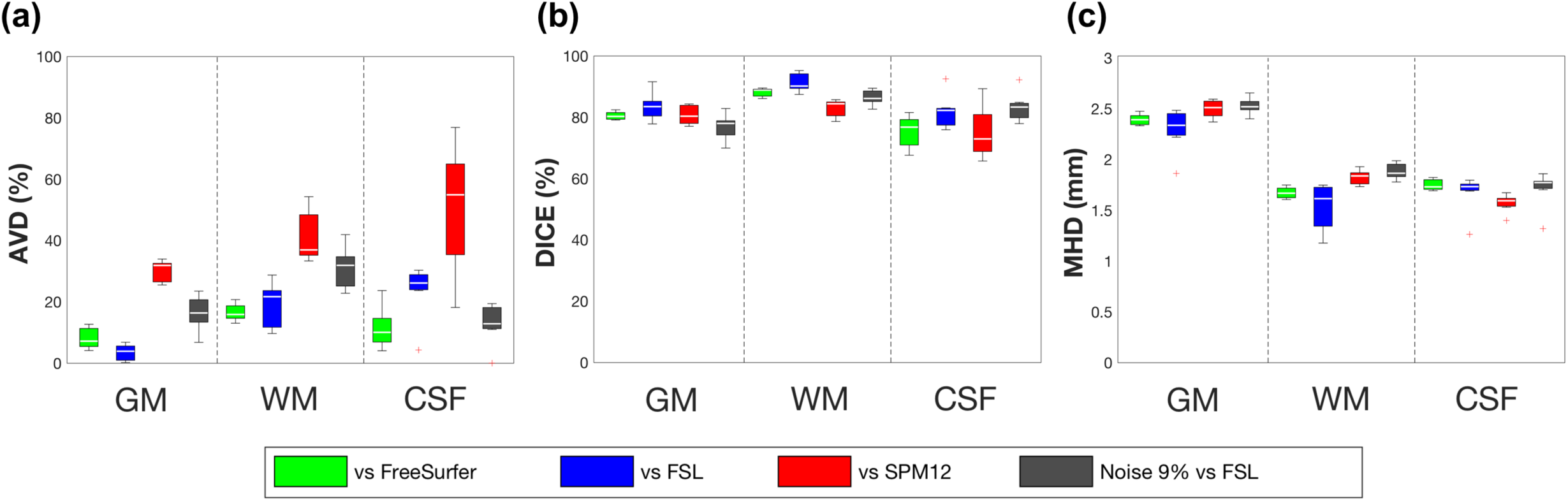
Similarity of segmentation with FreeSurfer, FSL, and SPM12. Box plots show AVD (a), DICE (b), and MHD (c) between the proposed method and the other methods. In addition, the segmentation at a noise level of 9% in images by the proposed method was evaluated with that by FSL at no noise. The red cross indicates outliers. AVD: absolute volume difference, DICE: dice coefficient, MHD: modified Hausdorff distance.

**Fig 6.**
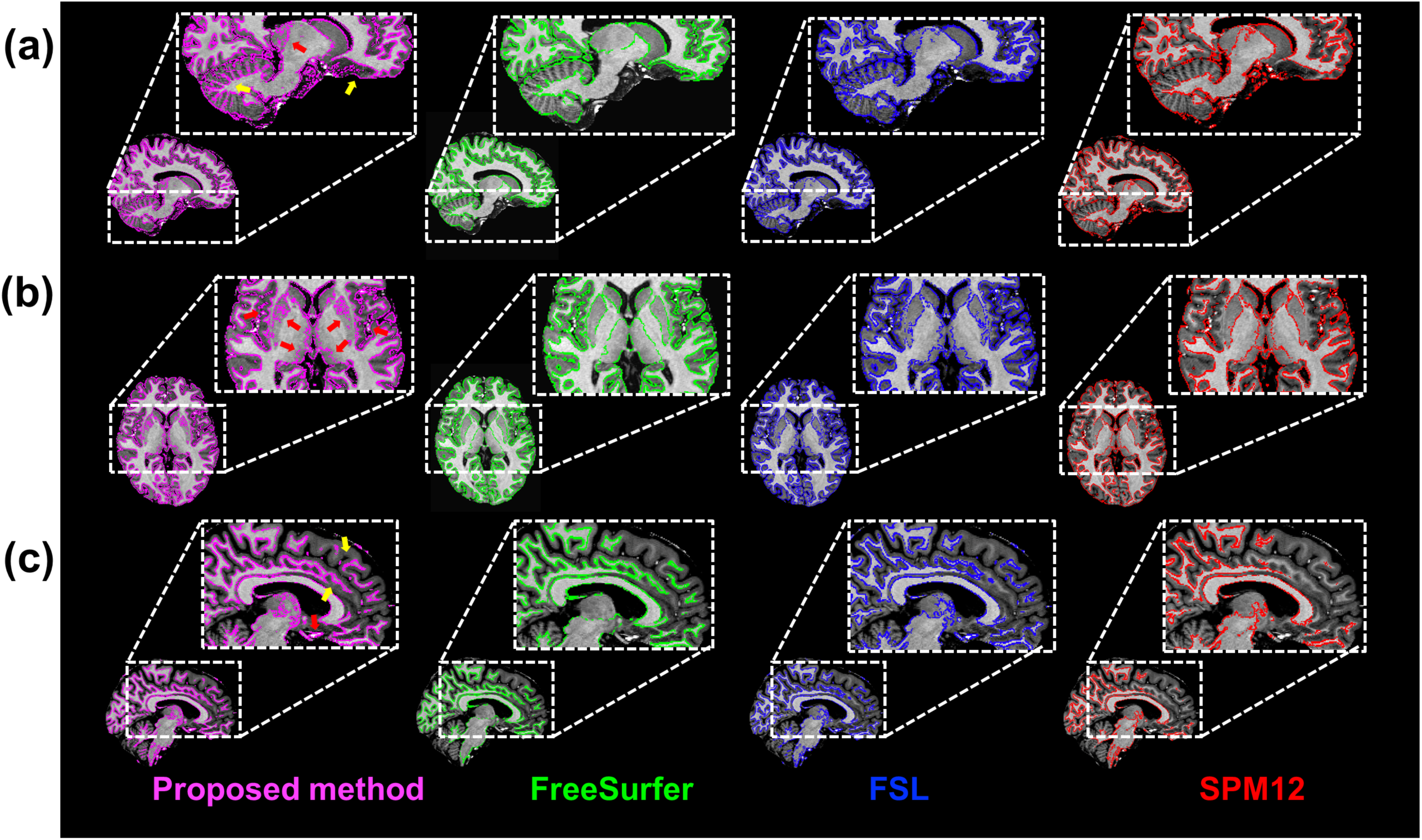
Segmentation performance using the proposed method, FreeSurfer, FSL, and SPM12. Gray matter (GM) segmentations in the medial part of the brain (a) and subcortical regions (b). White matter (WM) segmentation in the medial part of the brain (c). The yellow and red arrows indicate superior or inferior segmentation of GM and WM by the proposed method compared with the other methods, respectively.

## Discussion

We proposed a segmentation method with a significantly shorter processing time for brain tissues (GM, WM, and CSF). The proposed method employs a simple calculation of normalized signal intensities in the images with differing contrasts produced by an MP2RAGE sequence (UNI, INV1, and T1). The calculations were based on a consistent and specific tissue-dependent pattern of normalized intensities in the masks segmented by FreeSurfer, FSL, and SPM12, which are commonly used in neuroimaging protocols. Most segmentation methods use a single T1w image obtained from an MPRAGE sequence, and numerous groups have attempted to solve the overlapping signal distribution problem with complex and sophisticated methods that require considerable processing time to classify different brain tissues [9,11]. Although a few segmentation methods using multiple contrast images from an MP2RAGE sequence have been proposed [20], these require longer processing times because of the complexity of their algorithms. In contrast, the proposed method exhibits superior processing times because it utilizes simple calculations and three different contrast images from an MP2RAGE sequence. Therefore, the proposed method can segment a mask of brain tissues from the images, even with a high spatial-resolution at UHF, with shorter processing times.

To evaluate the segmentation performance by the proposed method, we calculated the AVD, DICE, and MHD values and compared them with other methods: FreeSurfer, FSL, and SPM12. We found differences between the proposed method and the other methods, but the brain tissue segmentation in the proposed method closely resembled that by FSL than those by FreeSurfer and SPM12. These differences in comparison with the other methods could be due to the differences in segmentation approaches in each method. Segmentation by the proposed method and FSL is based on the image intensities, whereas SPM12 and FreeSurfer use a template image as a reference for brain tissue and a parcellated brain atlas, respectively. The atlas-based segmentation approach can define subcortical structures, such as the putamen and caudate, and classify large blood vessels, resulting in its dissimilarity to the proposed method. In contrast, the GM and WM in the cerebellum and WM in the medial part of the brain areas showed good segmentation with the proposed method in comparison with the other methods, which also resulted in some of the comparative dissimilarities. Therefore, the dissimilarities with the other methods do not imply that the proposed method is inferior to the other methods because there is no gold standard software package for brain segmentation that performs as well as the manual method.

Our approach had some limitations as well as some potential for further development. One limitation was the poor segmentation of subcortical structures and large vessels. Two strategies can be implemented to overcome this limitation. First, because we employed different contrast images from an MP2RAGE sequence for segmentation and simple calculations, the performance of the proposed method was dependent on the relationship of the normalized intensities of the images. Therefore, the classification of the subcortical structure can be corrected using different relationships of normalized intensities with different acquisition parameters, e.g., TIs and FAs [16, 21, 22]. Second, compensation values can be employed in the segmentation calculation (i.e., thresholding), and these values can be optimized to improve the segmentation of GM and WM in subcortical structures and large vessels. Despite these limitations, the proposed method has the potential for use in segmentation because of the good segmentation of the cerebellum and WM in the medial part of the brain and the significantly reduced processing time. Although the boundary between GM and WM in the cerebellum is narrow and complex, due to the small size of its cytoarchitectural features [23], the cerebellum could be well segmented by the proposed method. Accurate segmentation of the cerebellum is useful for the reconstruction of an inflated image and the diagnosis of cerebellar WM diseases. The substantially shortened processing time is a critical factor because several studies have tried to reduce the processing time of segmentation for clinical diagnosis while maintaining acceptable accuracy [24] or allowing precise segmentation [25]. The proposed method can act as a quick reference for further, more precise brain segmentation approaches. The advantage of the shortened processing time will be more evident when using large datasets, such as those generated with ultra-high spatial resolution. In addition, although previous studies have suggested using both T1w and other MR images, such as PD, T2w, or fluid-attenuated inversion recovery images [6,26,27], for segmentation, these multi-spectrum segmentation approaches run the risk of head motion between scans. Because the proposed method uses different contrast images obtained simultaneously by an MP2RAGE sequence, it is free from the effects of possible head motion.

## Conclusions

We propose a novel brain tissue segmentation method that uses different contrast images from an MP2RAGE sequence. The proposed method allows rapid processing with relevant segmentation of brain tissues. A recent study has reported MP2RAGE-based segmentation using two images with different inversion times, i.e., INV1 and INV2 [28]. The study demonstrated superior segmentation in subcortical structures in comparison with FSL and SPM12, whereas our proposed method produced good segmentation in the cerebellum and WM in the medial part of the brain. Taken together, MP2RAGE-based segmentation is highly dependent on the relationship of the different contrasts between the input images used in the calculations, and this property could be of benefit in acquiring more precise focal segmentation of specific brain structures, such as the subcortical structures and the cerebellum. Thus, the MP2RAGE-based segmentation we proposed here has the potential to be applied, with optimized parameters of the MP2RAGE sequence, to both neuroimaging research and clinical diagnosis.

